# Inferring linguistic transmission between generations at the scale of individuals

**DOI:** 10.1101/441246

**Authors:** Valentin Thouzeau, Antonin Affholder, Philippe Mennecier, Paul Verdu, Frédéric Austerlitz

## Abstract

Historical linguistics strongly benefited from recent methodological advances inspired by phylogenetics. Nevertheless, no available method uses contemporaneous within-population linguistic diversity to reconstruct the history of human populations. Here, we developed an approach inspired from population genetics to perform historical linguistic inferences from linguistic data sampled at the individual scale, within a population. We built four within-population demographic models of linguistic transmission over generations, each differing by the number of teachers involved during the language acquisition and the relative roles of the teachers. We then compared the simulated data obtained with these models with real contemporaneous linguistic data sampled from Tajik speakers from Central Asia, an area known for its large within-population linguistic diversity, using approximate Bayesian computation methods. Under this statistical framework, we were able to select the models that best explained the data, and infer the best-fitting parameters under the selected models. This demonstrates the feasibility of using contemporaneous within-population linguistic diversity to infer historical features of human cultural evolution.

## 1. Introduction

Several recent studies use linguistic data under a computational framework aiming at reconstructing various aspects of the cultural history of human populations (Atkinson, 2011; Bouckaert et al., 2012; Gray and Atkinson, 2002; Pagel et al., 2013; Thouzeau et al, 2017). They rely on data mainly consisting in a set of presence or absence of linguistic items, within a given set of contemporaneous languages, which can be found, for example, in databases such as the World Atlas of Language Structures WALS (Dryer and Haspelmath, 2013), or the Global Database of Cultural, Linguistic and Environmental Diversity D-PLACE (Kirby et al., 2016). Thus, most studies consider languages at a macro-evolutionary scale, i.e. they deal only with differences among languages, neglecting the variability within each language. For instance, Gray and Atkinson (2002) used a set of Swadesh lists obtained for 87 languages to investigate the origin of the Indo-European linguistic family. Atkinson (2011) considered the number of phonemes used in 504 languages worldwide to test the hypothesis of a serial founder effect due to the Out-Of-Africa expansion. Reesink et al. (2009) used the linguistic diversity of the ancient Sahul continent (present-day Australia, New Guinea, and surrounding islands) among 121 languages to infer the history of the structural characteristics of these languages.

These approaches rely implicitly on several assumptions. They require primarily a clear separation between several differentiated languages. Nevertheless, this notion of distinct languages is often irrelevant at local scale, in particular in contexts of dialectal continuum or linguistic contacts (Heeringa and Nerbonne, 2001; Livingstone and Fyfe, 1999). Furthermore, most of these studies do not take into account within-population linguistic diversity, since traditional linguistics often considers languages as unique and coherent systems (Pateman, 1983). This assumption implies the loss of a large amount of information, knowing that the demographic phenomena occurring at population level – different population sizes, bottlenecks, expansions – are expected to play a major role in language evolution (Vogt, 2009). The inclusion of contemporaneous within-population linguistic diversity in the reconstruction of the demographic history of human populations at a local scale is thus expected to open a completely new dimension in the field of historical linguistic inferences.

In this context, Croft (1996) argued for the replacement of the ‘essentialist’ theory of language changes by a ‘population’ approach of these changes, and later proposed a detailed review of the “evolutionary linguistic” field and underlying paradigms (Croft, 2008). Nevertheless, very few previous studies dealt with the contemporaneous within-population linguistic diversity in a historical reconstruction perspective. Rodriguez-Larralde and Barrai (2000) used surnames of telephone users in Austria as linguistic contemporaneous information, showing that Austrian towns are subdivided into five main clusters with uniform levels of endogamy. Darlu et al. (2012) reviewed the analyses of paternally-inherited family names distributions, as an analogy to Y chromosomes, for historical inferences. Verdu et al. (2017) contrasted the proportion of African words in free speech among Cape Verdean Kriolu speakers with their proportion of African genetic admixture, showing that Cape Verdean genetic and linguistic admixture processes followed parallel histories, with possible co-transmission of genetic and linguistic variation. None of these studies developed an inferential approach that would allow researchers to distinguish among different historical mechanistic models, and inferring their constitutive parameters.

In order to perform such historical linguistic inferences from observed linguistic data, we need to assume one or several possible models of linguistic transmission between generations, and a possible set of historical scenarios that produced the observed data. Nevertheless, there is no consensus framework that allows handling within-population linguistic diversity data, in order to infer historical scenarios and evolutionary mechanisms. It requires to first build an explicit mechanism of linguistic evolution, and then to study the range of historical scenarios that could have produced the observed linguistic data. Nevertheless, the validity of the historical conclusions will depend on the validity of the assumed mechanism. It is, therefore, crucial to first determine the most relevant mechanism of linguistic evolution of a given set of linguistic objects, in order to produce, ultimately, valid inferences.

We evaluated here a series of models of linguistic evolution between generations at the individual scale. We did not study the history of higher-order objects such as “the languages”, but the history of the linguistic diversity carried by individuals within a population, among which communication events may occur over time. We aimed at understanding how the evolution of linguistic diversity among generations was affected by demographic parameters such as population sizes (the number of individuals of a given speech community), and thus to assess whether it was possible to infer the best demographic scenario and its corresponding parameters from a set of linguistic data.

Approximate Bayesian Computation methods (ABC, Beaumont et al., 2002; Tavaré et al., 1997) provide a particularly well-adapted framework to tackle this problem. In this paper, we used the recently developed Approximate Bayesian Computation via Random Forest (ABCRF) algorithm to assess, among a set of possible competing scenarios, the scenario that best explained the observed data, and to estimate the posterior parameters of this scenario (Breiman, 1999; Pudlo et al., 2016, Raynal et al., 2017).

For this purpose, we implemented an individual-based simulation program, which simulates the evolution of word variation among generations, under different modes of linguistic transmission. These simulated data allowed us to perform the ABCRF procedure on a real dataset from Central Asia. This dataset consisted of 30 individuals interviewed for 185 words across 10 villages in Tajikistan. These villages are known to use the same language, but with some variation across speakers (Mennecier et al., 2016). We aimed at inferring the most probable models of linguistic transmission mechanisms between linguistic generations, under a demographic scenario of population-size expansion or contraction.

We proposed four transmission models. The “*Clonal* model” assumed that each individual learns his/her linguistic words from only one teacher. The “*Sexual* model *1*” assumed that each individual learns his/her words from two teachers (one “male” and one “female”), with specific words transmitted only by males and others transmitted only by females. The “*Sexual* model *2*” assumed that each individual learned his/her words from two parents (one “male” and one “female”), without specific words belonging to males or females. The transmitting parent was drawn independently for each word, so that the set of words of an individual was a recombination between the sets of his/her two parents. Finally, the “*Social* model” assumed that each individual learns his/her words from the entire population. We aimed, then, at inferring, with ABC, the best-fitting parameters under the winning scenario: linguistic mutation rates, and population sizes. We demonstrated thus the feasibility of using contemporaneous within-population linguistic diversity to infer historical features in human cultural evolution.

## 2. Materials

We sampled cognate variation for 30 individuals from 10 Tajik villages in Central Asia (Figure 1) assuming that the individuals belonged to a single linguistic population. In contrast with our previous study, where we considered for each cognate only its most frequent variant in each locality (Thouzeau et al., 2017), we kept here the linguistic variants recorded for each individual separately. Thus, for each individual, we recorded the words used for 185 meanings from an adapted Swadesh list. Individuals were asked to state the most frequently used word for the associated meaning. We considered as “cognate” a group of words with the same etymological origin and the same meaning, such words being more likely to be related by a common ancestry (Atkinson et al., 2005). The phonetic transcriptions and classification of lexical data gathered in the field into cognates was performed by Philippe Mennecier similarly as in previous work (Mennecier et al., 2016; Thouzeau et al., 2017).

**Figure 1.**
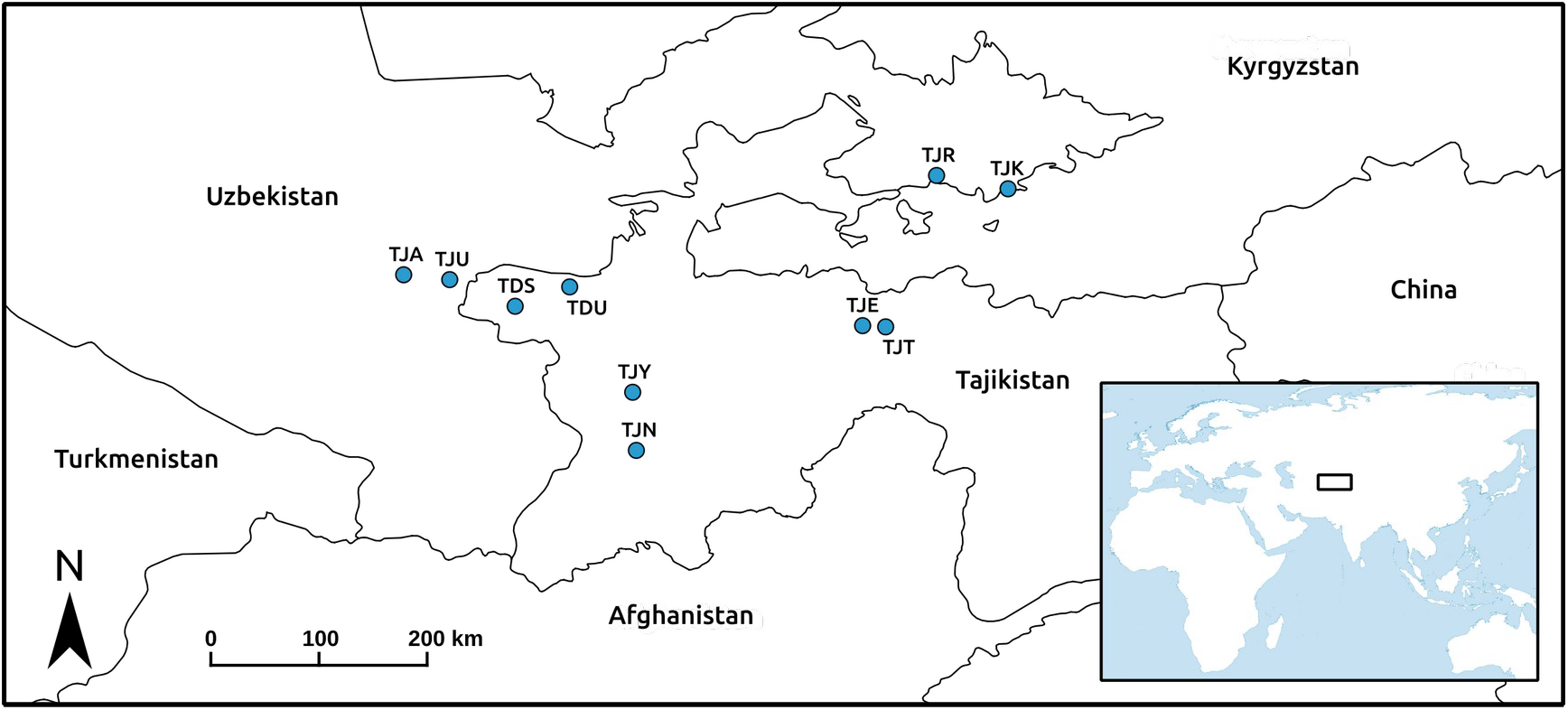
Geographical distribution of the 10 sampled units under study.

**Figure 2.**
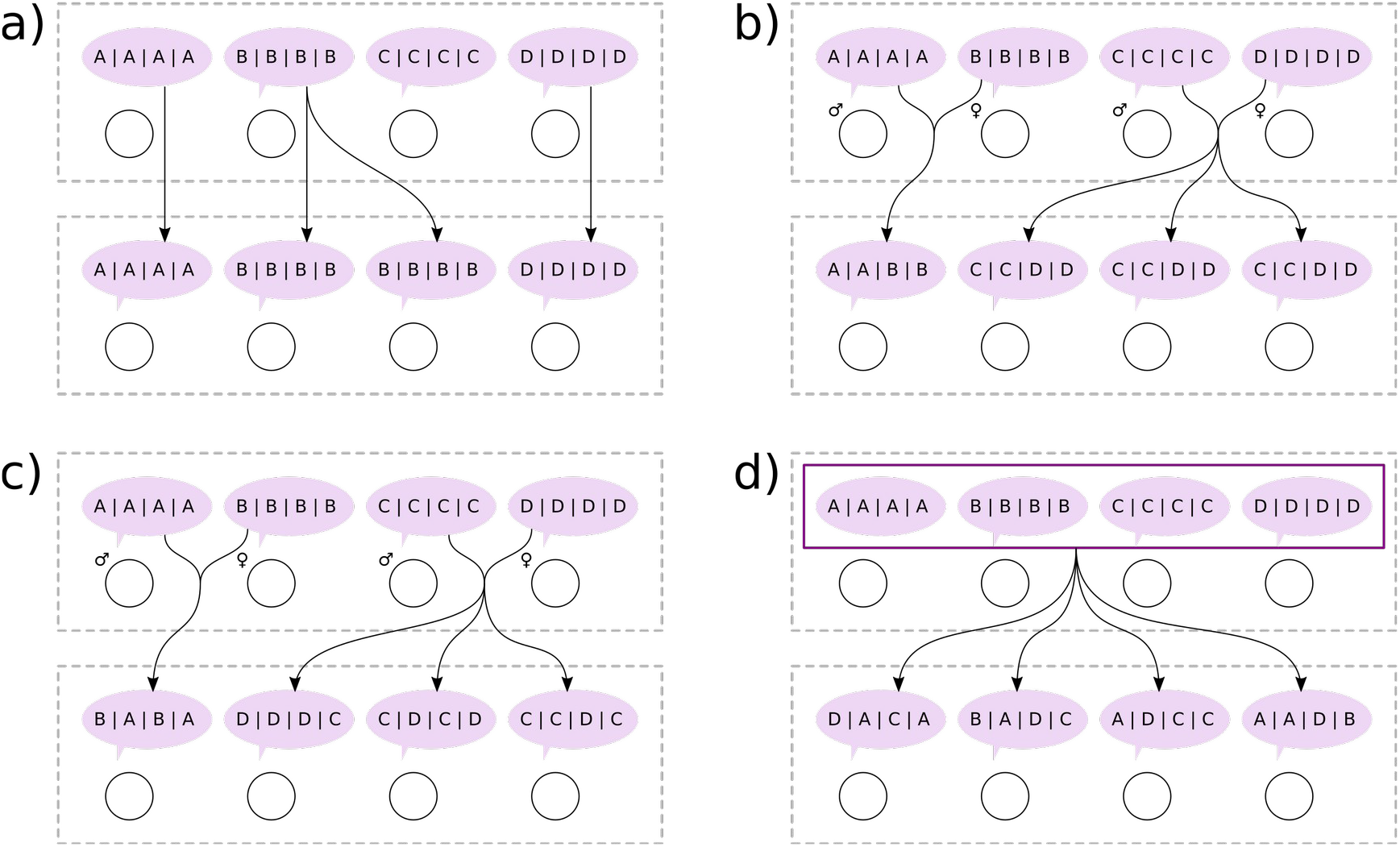
Four models of linguistic transmission between generations. Each circle represents an individual. The utterances that individuals produce depend only on the utterances that their teachers produced at the previous generation, and on the mutations induced during the transmission. Four transmission modalities were considered: (a) a “C*lonal*” model with only one teacher per learner, (b) a “S*exual 1*” model with two teachers associated with a distinct set of vocabulary for each sex, (c) a “S*exual 2*” model with two teachers without a distinct set of vocabulary for each sex, and (d) a “*Social*” model with the whole population as teacher for each learner.

## 3. Principle of ABCRF method

ABC methods were first introduced by Tavaré et. al. (1997) and Beaumont et al (2002), aiming to encompass the limitation of Markov chains Monte Carlo (MCMC) methods. For simple models, analytical formulas may be derived to compute the likelihood of the data under a given model. However, for complex models or for large datasets, computing the likelihood may be highly difficult and/or highly time consuming. ABC methods allow circumventing these problems by approaching the likelihood instead of exactly computing its value. It is a particularly well-suited statistical framework to develop within-population linguistic historical inferences tools, allowing to specify complex and explicit processes of linguistic interactions between a large set of agents.

ABC consists first in defining a set of models that could fit the observed data. Each model is characterized by several parameters, such as, but not limited to, effective population sizes and time of change in population sizes. A prior distribution for each parameter is chosen by the user, and corresponds to the range of values that are realistic for this parameter. A large number of simulations are then performed in order to generate data sets under the different models. For each simulation, each parameter of the model is drawn at random in its prior distribution. Sets of summary statistics are then computed on each simulated data set, each corresponding, therefore, to a given set of model parameters. In the ABCRF method (Pudlo et al. 2016), a random forest (RF) procedure is then applied to choose the best-fitting model. In short, the aim of the RF method is to produce a set of decision trees from the simulated data sets. Each tree is built by performing a supervised categorization of the whole set of simulations, according to the models which produced those simulations, and each one using a different subset of their summary statistics. Those subsets of summary statistics are selected randomly for each tree in order to improve classifications, because using the full set of summary statistics can lead to overfitting. The curse of dimensionality is also reduced by the RF procedure (Pudlo et al. 2016).

Then, the full set of summary statistics are computed on the real data, and this “observed” set of summary statistics is independently evaluated by each decision tree. Each one votes for a model, and the final decision is the majority of votes from the forest. Then, an error rate is computed to assess the confidence of this final decision. At this step, several models are usually rejected by the random forest.

For each selected model, another random forest is then constructed to estimate the parameters of the models. The principles of ABCRF regression are analogous to the principles of ABCRF classification (Raynal et al., 2017), but in this case, the trees use the summary statistics in order to predict the value of scalar transformed parameters. The forest is then built on the simulated summary statistics, in order to estimate the mean, median, and quantiles of the distributions of the real parameters.

## 4. Models

### 4.1. Production of utterances

We considered a linguistic population as a group of individuals that may potentially interact through linguistic communication. The mechanisms of linguistic communication and transmission may follow different modalities, which correspond to different models of linguistic evolution. In all cases, we considered that the unit of linguistic communication is the *utterance*, a production of words associated with a meaning (Croft, 1996).

We developed a general model of word transmission, which we applied in particular to the case of cognates, which correspond to words with different etymological origins that express the same meaning. For example, the Spanish word “Flor” and French word “Fleur” are two words with the same meaning (“Flower” in English) and the same etymological origin, and classified as the same cognate. The Spanish word “Mariposa” and French word “Papillon” are two words with the same meaning “Butterfly”, but with a different etymological origin. They are thus considered as different cognates. We considered here that cognates can vary among individuals within a population. This differs from the assumptions made in previous studies (Bouckaert et al., 2012; Gray et al., 2009; Thouzeau et al., 2017) where cognates are sampled at the language scale and for which individuals are considered as users rather than producers of the language.

### 4.2. Four models of acquisition of a new language

We developed a new Open Source C++ simulation software *PopLingSim 2* (*PLS2*, available at https://github.com/ValentinThouzeau/PLS2). This software implements an individual-based forward-in-time simulation model with discrete linguistic generations, in which we assumed that populations were composed of only two types of individuals: “learners” and “teachers”. The linguistic generation time corresponded to the time required for an individual between learning a language from teachers and teaching this language to learners at the following generation. We did not specify the linguistic generation time in our models, allowing it to be completely decorrelated from the reproductive generation time, and possibly much smaller. We assumed a neutral model in the sense that, even if the number of teachers per learner varied across models, the learners selected their teachers at random in the previous generation, with equal probabilities. We assumed that the rules of utterance productions of a teacher depended only on the utterances that he/she heard when he/she was a learner. We assumed that each learner chose only one word for each meaning during the learning phase. Two learners could choose the same word. After the whole learning phase, all teachers were discarded and all learners became teachers. Then, at the following generation, new learners appeared. The proportions of males and females were exactly 50%/50% at each generation. The models of linguistic acquisition differed by the number of teachers involved for each learner in his/her language acquisition process, and the relative roles of these teachers.

#### 4.2.1. Clonal Model

In the first model, named the “*Clonal*” model, each learner had only one teacher, which was drawn at random in the teacher population. The learner copied “in a clonal way” every word that the teacher produced. This would correspond, in genetics, to a clonal reproduction model, as observed e.g. for bacteria or for mitochondrial DNA and non-recombining regions of the Y chromosome in humans and other mammals, for instance.

#### 4.2.2. Sexual Model 1

In the second model, named the “*Sexual 1*” model, two different teachers (one “male” and one “female”) were drawn at random within the population for each learner respectively. The learner then copied directly the first half of the words produced by teacher 1, and the second half of the words produced by teacher 2. Thus, half of the words were always transmitted by one teacher, and the other half by the other teacher, the two different sets being always the same for all generations.

#### 4.2.3. Sexual Model 2

In the third model, named the “*Sexual 2*” model, two different teachers (one “male” and one “female”) were attributed to each learner at random. For each word, the learner copied at random either the word from teacher 1 or teacher 2, with equal probabilities (½, ½). Thus, no particular word had a teacher-specific transmission; every word was transmitted from one of the two teachers chosen at random. This is analogous, in genetics, to a sexual reproduction model with free recombination.

#### 4.2.4. Social Model

In the fourth model, named the “*Social*” model, for each meaning each learner copied a word drawn at random from all the words produced by all the teachers in the population. Thus, each learner learned his/her set of words randomly from the entire speech community, or rather, from all possible utterance variants of teachers for a given meaning at a given generation.

### 4.3. Mutation model

For each model, we assumed that errors could occur during the transmission of each word, leading to the creation of a completely new word. We denoted such errors “linguistic mutations”. The mean mutation rate per linguistic generation μ_L_ was drawn in a log-uniform prior distribution, between 10-6 and 10-1 mutations per word per generation. For each word, its mutation rate was subsequently drawn in a beta distribution with mean μ_L_ and shape parameter β = 2, allowing us to simulate a set of words each with different rates of change over time.

### 4.4. Historical scenario

We focused here on a single linguistic population, defined as the number of individuals that contributed significantly to the currently observed linguistic diversity, where the utterances of a sample of individuals have been obtained using a linguistic questionnaire in the final generation. This linguistic population evolved under a historical scenario (Figure 3), in which it was first of constant size *N*_0_ individuals involved in linguistic communications during *t*_0_ = 5×*N*_0_ generations. As we visually checked, this time was sufficient to reach an equilibrium between the production of linguistic diversity through mutation, and the reduction of this diversity through random sampling (i.e. linguistic drift). Then, this population underwent an instantaneous change of population size, reaching a new size *N*_1_, and the population remained at this size during *t*_1_ generations. This model allowed simulating a range of histories, depending on the relative values of the parameters *N*_0_ and *N*_1_ and on the value of *t_1_*. *N*_0_ and *N*_1_ were drawn in a uniform prior distribution bounded between 100 and 1000 individuals, this upper bound being set to limit the large computation-time requirements for completing forward-in-time simulations. These prior distributions reflected the uncertainty in the number of individuals that contributed significantly to the linguistic diversity observed in the sampled population. The size of this ancestral population was indeed completely unknown. AIndeed, even if some information could be obtained on the census size of the current population, it likely does not reflect the ancestral linguistic census sizes. Time *t*_1_ was drawn in a uniform prior distribution between 0 and 1000 generations. The median, the minimum, the maximum, and the 5% quantiles of the priors are summarized in Table 1.

**Figure 3.**
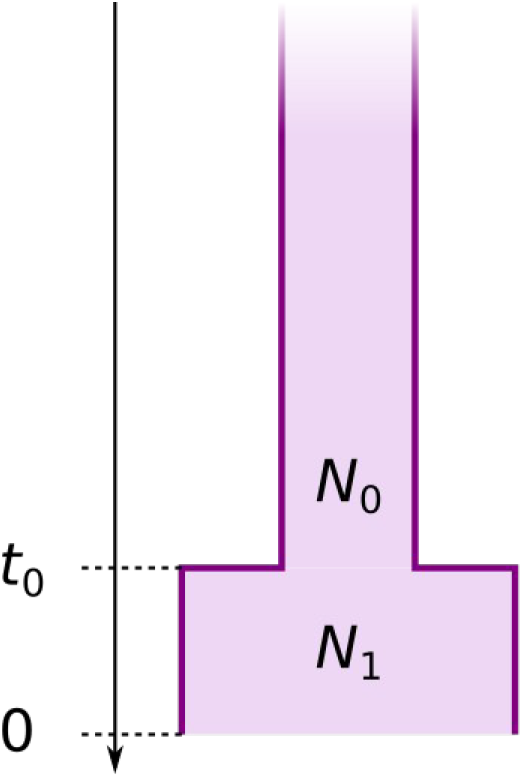
Historical scenario. If *N_0_* = *N_1_*, we assumed a scenario of constant population size. If *N_0_* < *N_1_*, we assumed a scenario of expansion of the population. If *N_0_* > *N_1_*, we assumed a scenario of contraction of the population.

**Table 1.** Summary of the prior distributions of the parameters for the four models.

|  | Median | Min | Max | Variance | Quantile 2.5% | Quantile 97.5% |
| --- | --- | --- | --- | --- | --- | --- |
| $N_0$ | 550 | 100 | 1000 | 67645 | 122 | 978 |
| $N_1$ | 550 | 100 | 1000 | 67645 | 122 | 978 |
| $t_1$ | 500 | 0 | 1000 | 83490 | 25 | 975 |
| $\mu_L$ | $3.165 \times 10^{-4}$ | $10^{-6}$ | $10^{-1}$ | $3.58 \times 10^{-4}$ | $1.35 \times 10^{-6}$ | $7.73 \times 10^{-2}$ |
| $N_0 \times \mu_L$ | 0.150 | $10^{-4}$ | 100 | 141.91 | $5.25 \times 10^{-4}$ | 44.5 |
| $N_1 \times \mu_L$ | 0.150 | $10^{-4}$ | 100 | 139.05 | $5.25 \times 10^{-4}$ | 44.5 |
| $t_1 \times \mu_L$ | 0.116 | 0 | 100 | 129.55 | $2.80 \times 10^{-4}$ | 42.0 |

## 5. Analyses

### 5.1. Simulations

For each model, we performed 10 000 simulations using our newly-developed software *PopLingSim 2* (*PLS2*). We parallelized the simulations using 250 cores of the cluster station *Genotoul*, amounting to approximately 90 000 CPU hours. Most of this computation time was spent during the phase to reach equilibrium between mutation and drift at *t_0_* = 5×*N*_0_ generations.

During the process of sampling words from our simulations, we simulated missing values by transforming cognates drawn at random into missing values; the total number of simulated missing values was set to the number of missing values in the real data set, to avoid the bias they may induce in the following ABC procedures.

### 5.2. Summary statistics

We constructed a new set of population linguistic summary statistics. We computed *p_i,j_*, the proportion of individuals using the word *i* of the meaning *j*, and then computed the linguistic diversity 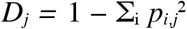 analogous to genetic diversity (Nei, 1987). We also computed chi-square values, over 200 pairs of randomly sampled meanings: 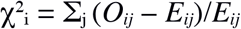. The observed value *O_ij_* corresponds to the number of individuals for which a pair of words *j* is observed for meaning *i*. The expected value *E_ij_* corresponds to the number of individuals for which this pair of words *j* would be observed for meaning *i* if the words were randomly distributed among individuals. We computed correlation coefficients values, over 200 pairs of meanings randomly sampled, as *r* = (*p_i,i’,j,j’_* − *p_i,j_ p_i’,j’_*) / [ *p_i,j_* (1 − *p_i,j_*) *p_i’,j’_* (1 − *p_i’,j’_*)]^−1/2^, with *p_i,i’,j,j_* the proportion of pairs of individuals using the word *i* of the meaning *j* and the word *i’* of the meaning *j’*. We then computed the frequency spectrum of the number *i* of words per meaning, *F_i_*.

Then, we computed across all words:

- The mean linguistic diversity,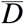;
- The range of linguistic diversity, R(*D*);
- The variance of linguistic diversity, V(*D*);
- The mean number of words per meaning, 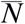;
- The variance of the number of words per meaning, V(*N*);
- The mean number of different words between two individuals,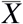;
- The range of the number of different words between two individuals, R(*X*);
- The variance of the number of different words between two individuals, V(*X*);
- The mean of the chi-square values,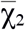;
- The range of the chi-square values, R(χ_2_);
- The variance of the chi-square values, V(χ_2_);
- The mean of the correlation coefficients values, 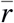;
- The range of the correlation coefficients values, R(*r*);
- The variance of the correlation coefficients values, V(*r*);
- The minimum of the frequency spectrum, min(*F*)
- The maximum of the frequency spectrum, max(*F*)
- The mean of the frequency spectrum, 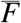
- The mode of the frequency spectrum, mode(*F*)
- The range of the frequency spectrum, R(*F*)
- The 25th quartile of the frequency spectrum, *F_25_*
- The median of the frequency spectrum, *F_50_*
- The 75th quartile of the frequency spectrum, *F_75_*
- The three axis of the linear discriminant analysis of the previous statistics, LD_1_, LD_2_, LD_3_.

### 5.3. Power analysis on simulated data

We performed a power analysis of the model selection procedure, to evaluate the impact of the number of sampled individuals, the number of sampled words, and the number of simulations on the prior error rate, i.e. the number of cases in which a wrong model was selected among the four possible models by the ABCRF model-choice procedure. This was done for a total of 61 situations in which we varied the number of sampled individuals between 2 and 100, the number of sampled words per individuals between 2 and 300, the number of simulations between 1,000 and 10,000. This maximal value of 10,000 simulations was due to the high computational cost of forward-in-time simulations. In each case, we computed the prior error rate through cross-validation using the function *abcrf* of the R package *abcrf*. This procedure considers, in turn, each simulation under each competing model as a pseudo-observed data, and performs the ABCRF model-choice using all other simulations in the reference table for training the random forest.

### 5.4. Model selection on real data

Before model selection, we performed a goodness-of-fit test to check whether the simulations were able to produce data close to the real data using the function *gfit* from the *R* package *abc* (Csilléry et al., 2012). We performed model selection using the *R* package *abcrf* with the RF algorithm and the function *abcrf* (Pudlo et al., 2016). We graphically checked if a forest of 500 trees allowed a convergence of the error rate. We computed the variables importance, indicating which variables have the most predictive power. We also performed a cross-validation analysis using an out-of-bag approach implemented in the function *abcrf* of the package *abcrf*, evaluating how the algorithm *a priori* distinguished between the four models.

For the selected model, we then selected the 100 simulations which were closest to the real data, based on the Euclidean distance of the statistics that were standardized for a mean of 0 and a variance of 1. We then tested whether the random forest algorithm was able, in this region of simulated data close to the real data, to correctly select the true model.

### 5.5. Parameters estimation on real data

We used the RF algorithm with the function *regAbcrf* of the package *abcrf* (Raynal et al., 2017) to estimate the expectation, median, variance and 5% quantiles of the parameters *N*_1_, *N*_0_, *t*_1_, μ_L_ and of the composite-parameters *N*_1_×μ_L_, *N*_0_×μ_L_ and *t*_1_×μ_L_. Note that the RF algorithm does not estimate the entire posterior distribution of the parameters directly, but estimates the quantiles of this distribution instead.

## 6. Results

### 6.1 Power analysis

Using simulated data under the four competing linguistic transmission models, we showed that an increase in the number of words sampled beyond 185 words increased moderately the power of the analyses (Figure 4). We found also that the decrease in error followed an exponential decay profile (Figure S1). Increasing the words sampling effort by several orders of magnitude would therefore be necessary to significantly reduce model selection error. Increasing the number of sampled individuals beyond 30 individuals increased only slightly the statistical power of the analysis (Figure 5), which converged towards a limit value. An increased sampling effort on the number of individuals could also, therefore, only moderately reduce the model selection error. Finally, we showed that the model-selection prior error rate converged with 10000 simulations (Figure 6), which indicated that increasing the number of simulations could not lead to a lower error.

**Figure 4.**
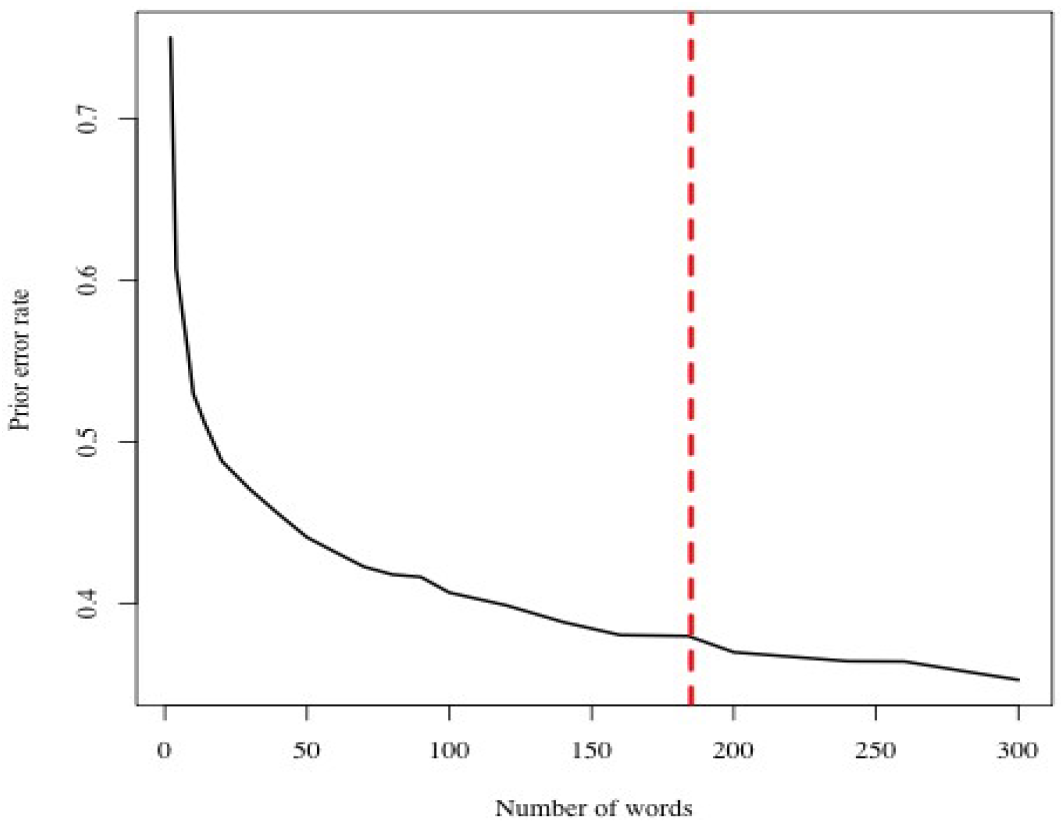
Prior error rate depending on the simulated number of sampled words, with 30 sampled individuals and 10000 simulations. The red dashed line indicates the number of words of the real sample.

**Figure 5.**
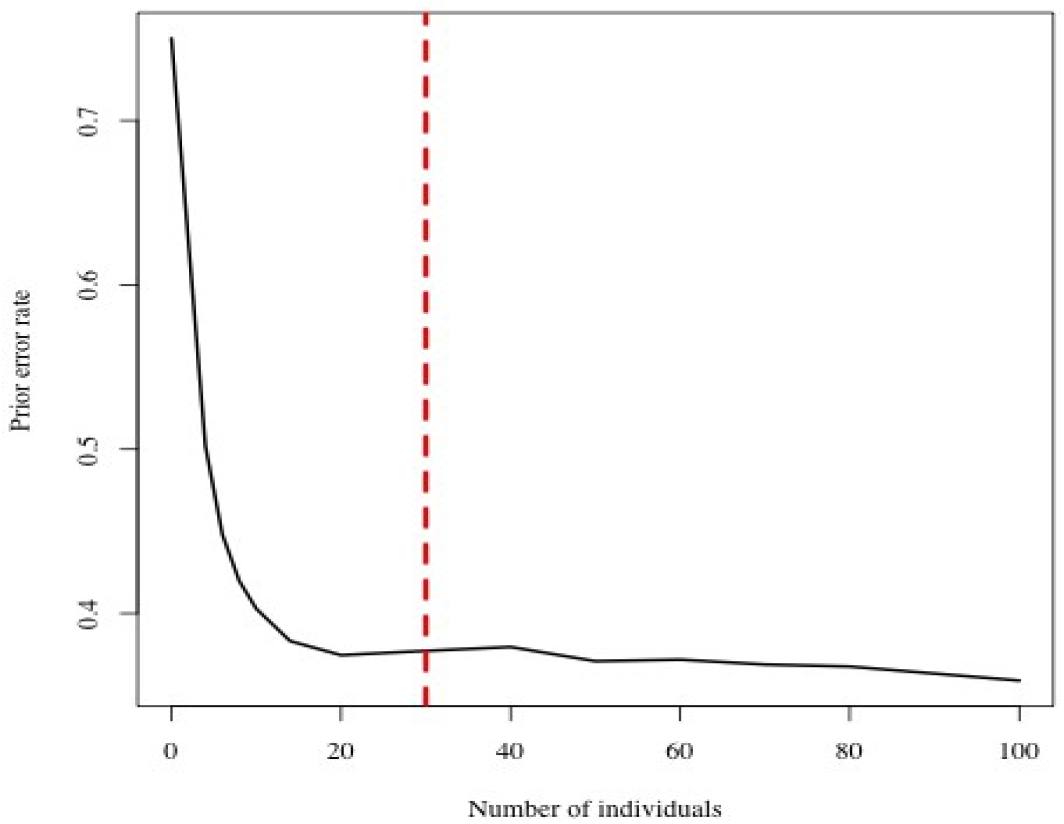
Prior error rate depending on the simulated number of sampled individuals, with 185 sampled words and 10000 simulations. The red dashed line indicates the number of individuals of the real sample.

**Figure 6.**
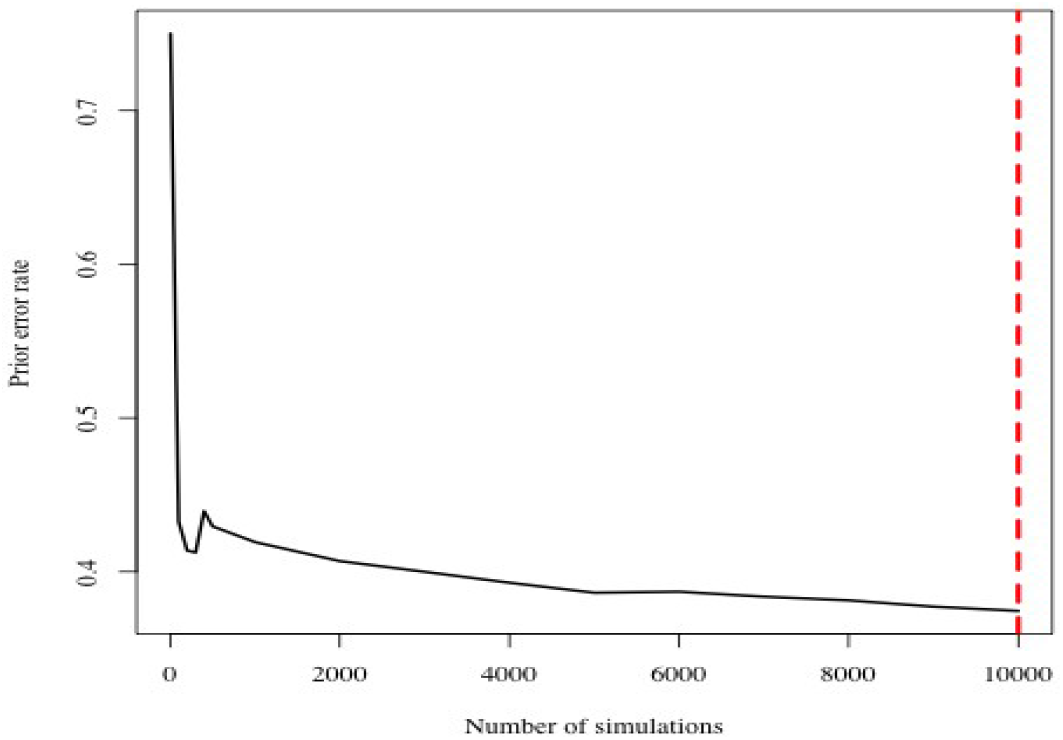
Prior error rate depending on the number of simulations, with 30 sampled individuals and 185 sampled words. The red dashed line indicates the value used for the analyses.

### 6.2. Model selection

Using the goodness-of-fit test, we verified that there was no significant differences between the real and simulated datasets (*p*-value = 0.71, with 1000 replications). We performed the RF analysis using 500 trees, and verified graphically that the error rate converged. The number of trees voting for the second model was 487 out of 500 (Table 2). The RF analysis thus rejected the *Clonal*, *Sexual 2* and *Social* models, and selected the *Sexual 1* model for the real data with a posterior probability *p* = 94.4%.

**Table 2.** Proportion of votes for the four models of linguistic evolution.

| Clonal | Sexual 1 | Sexual 2 | Social |
| --- | --- | --- | --- |
| 11 | 487 | 1 | 1 |

The variable importance analysis showed that the main statistics used by the RF procedure to select the models were the first two axes of the linear discriminant analysis LD_1_ and LD_2_, the variance of the number of different words between two individuals V(*X*), and the variance of the correlation coefficients values V(*r*) (Figure S2).

The cross-validation analysis on simulated datasets (Table 3) indicated a good *a priori* differentiation between the *Clonal* model and other models, with about 76% of simulated datasets under this clonal model correctly assigned to the true model. Similarly, the *Sexual 1* model was correctly attributed for about 76% of the simulated datasets. On the other hand, the *Sexual 2* model and the *Social* model were difficult to distinguish *a priori*, as the simulated datasets from these two models were arbitrarily attributed to one or the other by the cross-validation procedure.

**Table 3.** Confusion matrices from the out-of-bag cross-validation analysis of the four models, using 10 000 pseudo-observed data.

|  |  | True model |  |  |  |
| --- | --- | --- | --- | --- | --- |
|  |  | Clonal | Sexual 1 | Sexual 2 | Social |
| Selected model | Clonal | 7620 | 1785 | 282 | 313 |
|  | Sexual 1 | 1358 | 7698 | 439 | 505 |
|  | Sexual 2 | 283 | 816 | 4782 | 4119 |
|  | Social | 276 | 805 | 4200 | 4719 |

The RF algorithm assigned to the correct model 100% of the simulations produced by the *Sexual 1* model which were closest to the real data. Compared to the global cross-validation results, this indicated that the method performed better in selecting the correct model in the region of the parameter space occupied by the real data, than in the entire space occupied by simulations.

### 6.3. Parameter estimation

For the selected model (*Sexual 1*), we could estimate the linguistic mutation rate (μ_L_) on the real data: the quantiles of its posterior distribution were much narrower than the quantiles from its prior (Table 4). We estimated that this rate ranged between 1.61×10^−4^ and 1.50×10^−3^ mutations per cognate per linguistic generation at the 95% credibility level. Conversely, we could not estimate the demographic parameters (*N*_1_, *N*_0_, and *t*_1_), for which posterior quantiles did not differ substantially from prior quantiles. However, we could estimate the composite parameters *N*_1×_μ_L_ *N*_0×_μ_L_ and *t*_1_μ_L_, for which posterior quantiles were substantially narrower than those of their respective priors. There was no clear evidence of expansion or contraction, since the confidence intervals of *N*_1×_μ_L_ and *N*_0×_μ_L_ overlapped.

**Table 4.** Summary of the posterior distributions of the parameters, assuming a *Sexual 1* scenario.

|  | Expectation | Median | Variance | Quantile 2.5% | Quantile 97.5% |
| --- | --- | --- | --- | --- | --- |
| $N_0$ | 489 | 523 | 70157 | 112 | 928 |
| $N_1$ | 541 | 547 | 69571 | 105 | 949 |
| $t_0$ | 548 | 541 | 99439 | 48 | 985 |
| $\mu_L$ | $5.42 \times 10^{-4}$ | $4.14 \times 10^{-4}$ | $1.8 \times 10^{-7}$ | $1.61 \times 10^{-4}$ | $1.50 \times 10^{-3}$ |
| $N_0 \times \mu_L$ | 0.250 | 0.139 | 0.081 | 0.036 | 0.967 |
| $N_1 \times \mu_L$ | 0.178 | 0.155 | $4.35 \times 10^{-3}$ | 0.098 | 0.347 |
| $t_1 \times \mu_L$ | 0.358 | 0.215 | 0.148 | 0.010 | 1.15 |

## 7. Discussion

We built here individual-based models simulating the linguistic evolution of a population, under given demographic scenarios, considering four possible types of linguistic transmission between generations. We used an ABCRF framework (Pudlo et al, 2016, Raynal et al, 2019) to compare the simulated data with a real dataset of 30 individuals in Central Asia typed for 185 words, in order to estimate which model fitted best the data and estimate the parameters of the selected model.

ABC relies on approximating the likelihood of the data by that of a set of summary statistics, *a priori* informative about the historical process to be inferred. ABC was initially developed with summary statistics explicitly linked to the parameters of interests, and therefore highly informative for accurate ABC inference (Tavaré 1997). However, for most case studies, it is not known, *a priori*, which summary statistics will be informative for ABC inference (Blum et al. 2013). Several complex statistical approaches have been developed, therefore, to select *a priori* sets of relevant summary statistics for ABC inference and to overcome the dimensionality curse and parameter posterior identifiability issues, which result from considering very large numbers of summary statistics, possibly correlated and unevenly informative (Csilléry et al. 2012; Blum et al. 2013; Prangle 2015). Importantly, ABCRF model choice inference is unaffected by the dimensionality curse faced by most other ABC model-choice frameworks, as each decision tree is built with random subsets of summary statistics (Pudlo et al. 2016). However, the accuracy of model parameter inference in ABC, whether using RF or another approach, still relies on finding minimal subsets of highly informative summary statistics (Raynal et al. 2017), which therefore requires empirical case-by-case testing of novel sets of summary statistics.

A main advantage of the ABC framework is its high flexibility, which will allow researchers, in future work, to include more sophisticated models with additional parameters of interests to linguistic evolution. Moreover, ABC offers a model selection procedure that has no equivalent under an analytic framework, and it offers also the possibility to compute the credibility interval of the inferred parameters, which would require a fully stochastic approach in an analytical framework. We showed, first, that some of our models were able to produce simulated data close to the contemporaneously observed data. Therefore, our approach implements realistic individual-based linguistic transmission, consistent with the observed linguistic diversity of the sampled populations.

We also provided inferences on some features of linguistic history of Tajik-speaking individuals, selecting the most plausible mechanisms of linguistic transmission among the competing options tested, and estimating the parameters of the selected models for our sample. The high posterior probabilities of the S*exual* model 1, compared to the other models, indicated that the mechanisms of linguistic acquisition followed, in this study-case, a process of linguistic transmission from two teachers with their own vocabulary. In other words, we inferred that these individuals did not learn their basic vocabulary from only one individual, nor from two individuals without “sex”-specific vocabulary, nor from the whole speech community. We estimated that they learn their vocabulary from two individuals with “male”-specific and “female”-specific words. This linguistic-transmission mechanism may reflect the fact that Tajik populations are cognatic (Krader, 1966), i.e. they inherit social status and material goods from their two parents. This symmetric role of parents in cultural transmission across generations appears thus to be reflected linguistically, as learners appear to receive specific words from both parents. Future studies on populations with other lineage and kinship descent systems, such as patrilineal or matrilineal descent rules, will allow better understanding how social-descent rules and features may influence linguistic transmission processes in a given population.

Our estimates of the mean linguistic mutation rate of the words of the Swadesh list in this population ranged between 10^−4^ and 10^−3^ mutations per word per generation. Interestingly, the mutation rate estimated here fell in the same range as the mutation rate estimated in previous macro-evolutionary linguistic studies (Pagel et al., 2007). Considering that languages at a global scale emerge from the interactions among individuals, we may thus hypothesise that the mutation rate estimated globally emerges from the mutation rate at a local scale. Under this assumption, further studies could investigate whether macro-evolutionary linguistic processes (i.e. processes occurring at the scale of a whole language or a linguistic variety), may also emerge from micro-evolutionary linguistic processes (i.e. at the scale of the individuals within a population).

Population genetics effective population sizes estimated in Central Asia differ from census population sizes (Palstra et al. 2012). Similarly, the estimated linguistic population size in our model did not necessarily reflect the real size of the community, and effective linguistic populations are possibly much smaller in size than empirical groups of speakers. Our posterior estimates of the number individuals that contributed significantly to the observed linguistic diversity did not differ from the priors of the simulations. It meant that our method could not directly estimate the number of individuals in the current and ancestral linguistic populations, but only synthetic parameters such as *N*_0*μ*_. In this context, a perspective might be to design specific summary statistic to improve our ability to infer the number of individuals that contributed significantly to the observed linguistic diversity. Another promising approach might be to sample individuals in the population at different moments in time, separated by at least several decades, analogously to what is done in population genetics, where it is the most efficient method for estimating recent population sizes, independently of mutation rate (Foll et al., 2014).

In this study, unlike most other studies focusing on within-population linguistic diversity (Baxter et al., 2009; Danescu-Niculescu-Mizil et al., 2013; Kandler et al., 2010), we only used contemporaneous linguistic diversity. This method allowed us to perform historical inferences based only on sampling campaigns conducted in existing populations. The amount of available information depended only on the sampling effort, and not on the availability of dense historical records, which are unavailable for numerous languages. It would be of great interest in future works to be able to distinguish among the S*exual 2* model (with only two teachers) and the *Social* model (with a whole community as a teacher). As we showed in the power analysis, increasing the sampling effort (in terms of number of individuals or in term of number of words) was not sufficient to reliably distinguish between these two models, using our set of summary statistics. As for the inference of demographic parameters, developing new summary statistics and/or designing multi-generational studies might be the best solution to further distinguish among closely related linguistic transmission modes in future work.

Our approach could be extended in several other ways. First, the linguistic acquisition models that we proposed here did not integrate the particular constraints of communication processes. In particular, we assumed a neutral production of variants without any constraints on linguistic communication. Some evolutionary linguists would argue for an integration of the particularity of languages as communication systems, associated with a strong set of constraints (Beckner et al., 2009). Indeed, individuals maximize their probability of being understood, while minimizing their communication costs, two features that strongly affect linguistic evolutionary processes (Tamariz and Kirby, 2015). These constraints are particularly strong in the case of phonological, morphological, or syntactical systems, and we may wonder whether lexical variants are also subjected to these constraints. If so, these particularities of linguistic systems may be at odds with inferences based on a model of neutral evolution, and should thus be taken into account for a more accurate model of linguistic evolution at the individual scale, for historical inferences purposes.

It will also be of interest to study the transmission model of other types of linguistic objects, for instance focusing on other types of words such as food lexica, or very recently acquired technological lexica. Those different types of words could be transmitted differently, and our results could be different in the case of these particular lexical elements. Other types of linguistic data could also be obtained, like phonetic productions or syntactic rules, and it could be then assessed whether these linguistic elements are transmitted or not in the same way as the words of the Swadesh list. In addition, individuals may know more variants than those they use most frequently. It may then be possible in future works to model also the evolution of word usage, in order to take into account a greater part of the lexical diversity of languages.

We assumed also that linguistic transmission occurs between two linguistic generations, ignoring the impact of communications between individuals of the same generation. Moreover, we did not take into account global media such as books, radio, internet, or television. We will thus consider in further investigations several alternative models of language evolution, where the acquisition of language results from a series of interactions between individuals, who would update their language during each conversation.

Finally, note that the formalism of our models are close to the formalism of population genetics. This should allow us to develop joint inferences coupling genetic and linguistic data for the same set of populations and individuals. However, some theoretical limits remain. We may wonder whether a speech community (a “linguistic population”) is identical to a reproductive group (a “genetic population”). It is far from obvious that human reproductive boundaries overlap language boundaries among human groups. A joint model between genetics and linguistics would thus require clarifying and articulating rigorously the concepts of population genetics with the concepts of population linguistics to propose robust joint inferences.

## 8. Acknowledgements

We thank the Genotoul bioinformatics platform (Toulouse, Midi-Pyrénées) for providing help, computing and storage resources. V.T. was financed by a PhD grant from the French ‘Ministère de l’Education Nationale, de l’Enseignement Supérieur et de la Recherche’. V.T. and F.A. received a travel grant from the NEFREX project funded by the European Union (People Marie Curie Actions, International Research Staff Exchange Scheme, call FP7-PEOPLE-2012-IRISES). This work was also partially funded by the Agence Nationale de la Recherche grant DemoChips (ANR-12-BSV7-0012).

---

**Figure S1.**
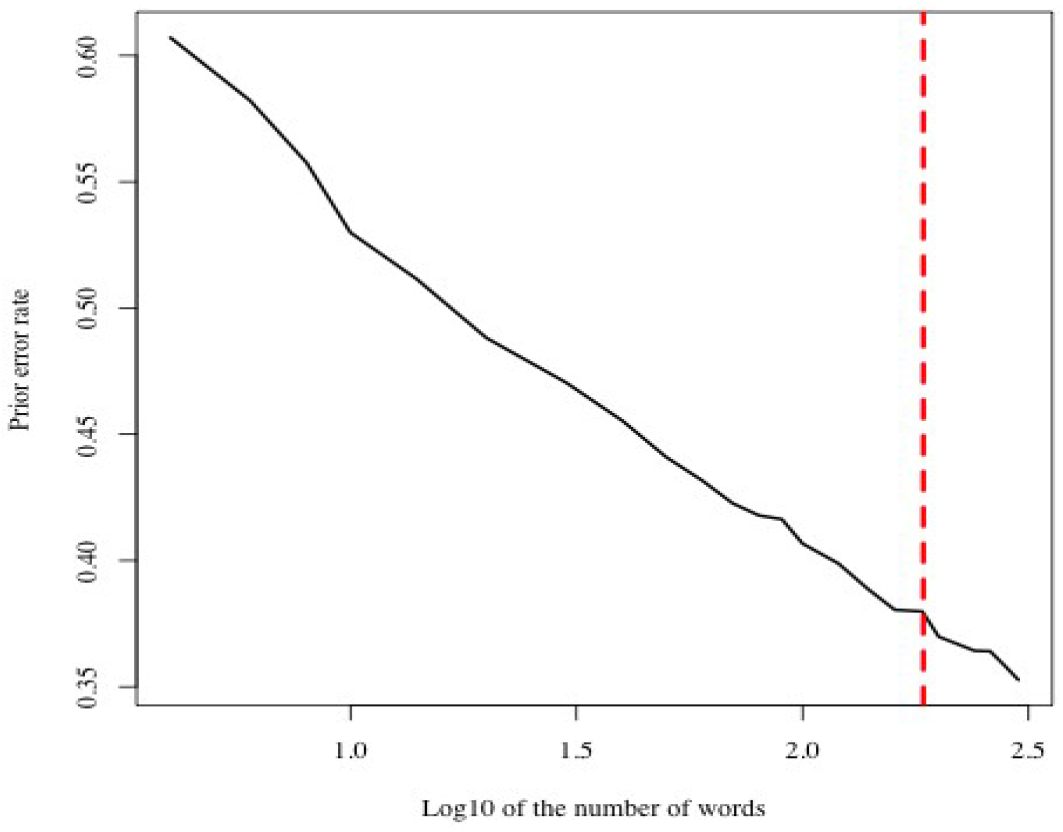
Prior error rate depending on the simulated decimal logarithm of the number of sampled words, with 30 sampled individuals and 10000 simulations. The red dashed line indicates the number of words of the real sample.

**Figure S2.**
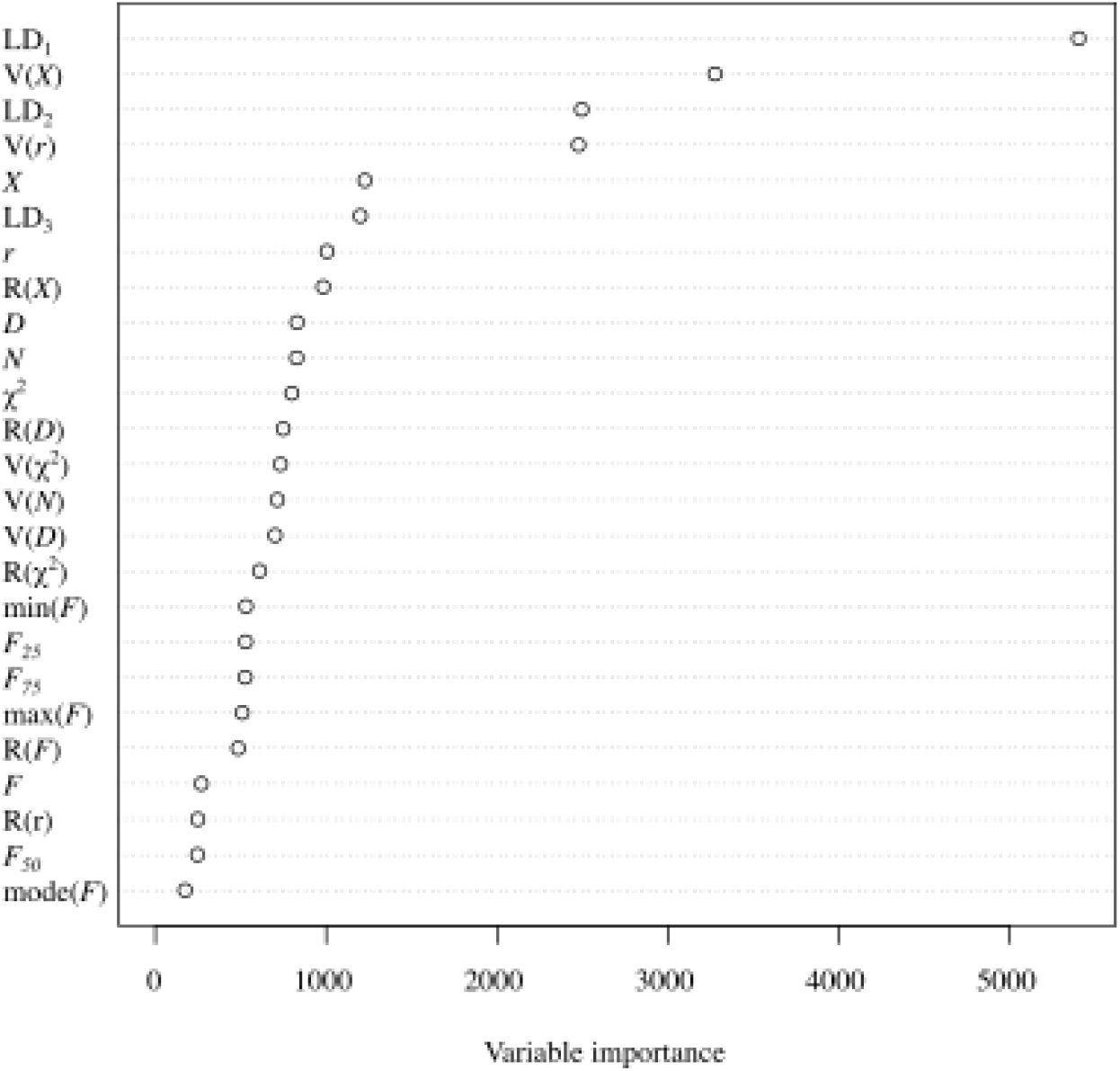
Variable importance of the random forest built for the model selection.

